# Comparative Evaluation of Commercial Iron Oxide Particles for Magnetic Particle Imaging Using Relaxometry and Image-Based Metrics

**DOI:** 10.64898/2026.09.15.751874

**Authors:** Bijita Neupane, Paula J Foster

## Abstract

In magnetic particle imaging (MPI), signal is generated from the non-linear magnetic response of superparamagnetic iron oxide (SPIO) nanoparticles. The optimization of SPIOs for MPI is an active area of investigation. Tracer performance can be evaluated by magnetic particle relaxometry (MPR) and image-based metrics. This study characterized and compared the performance of ten commercially available tracers, spanning a broad range of magnetic core sizes, hydrodynamic diameters and surface coatings, using the peak signal intensity measured from MPR, and the total signal and maximum signal intensity measured from 2D MPI images. Synomag-D tracers exhibited the highest MPR peak signal intensities and highest maximum signal per µg iron. ProMag, a micron-sized iron oxide particle (MPIO), yielded a low MPR peak signal intensity but had the highest total MPI signal per µg of iron. A strong correlation was observed between MPR peak signal intensity and MPI maximum signal intensity (R^2^=0.94). In contrast, there was a weak correlation between MPR peak signal intensity and MPI total signal intensity (R^2^=0.38), though this correlation improved when MPIOs were excluded from the analysis (R^2^=0.91). While MPR generally predicts the tracer performance, it does not completely replicate the complex imaging conditions. As such, comprehensive tracer evaluation requires combining both MPR and image-based metrics.

## INTRODUCTION

Magnetic particle imaging (MPI) is a relatively new imaging modality that directly detects superparamagnetic iron oxide (SPIO) particles. The basic principles of MPI are described briefly here and additional details can be found in comprehensive reviews [1–3]. In MPI, a set of opposing gradient coils create a selection field (gradient field) with a field-free region (FFR) near the centre. Particles outside of the FFR are magnetically saturated by the selection field, while particles within in the FFR remain sensitive to the excitation field (drive field), an alternating low amplitude magnetic field. The MPI signal is generated by the nonlinear response of SPIO particles to the excitation field. This response is not instantaneous and is governed by two primary relaxation mechanisms: Neel relaxation, which involves the internal rotation of the magnetic moment of the particle, and Brownian relaxation, which involves the physical rotation of the magnetic particle to align with the magnetic field.

MPI sensitivity is strongly influenced by the magnetic properties of the SPIO tracer. Magnetic particle relaxometry (MPR) is commonly used to measure the intrinsic particle sensitivity when developing or screening SPIOs for MPI. MPR measures the magnetization response of a sample of SPIO directly, in a controlled localized field environment, and the output is a point spread function (PSF). Particle sensitivity is reported as the peak amplitude of the PSF normalized to iron mass of the sample. Imaging can also be used to evaluate tracer sensitivity. This is usually done using a calibration line generated by imaging a number of samples containing known amounts of iron. The maximum signal intensity in the region of interest (ROI) or the total MPI signal (mean signal x ROI size) for each sample is plotted against iron mass [4, 5]. The slope of the resulting calibration line reflects the amount of MPI signal generated per unit iron. A steeper slope indicates that smaller amounts of iron produce larger MPI signals, corresponding to improved tracer detectability. While MPR measures the maximum potential performance of a tracer, imaging evaluates its functional performance within a complete MPI system incorporating not only the magnetic properties of tracers, but also scanner hardware, acquisition parameters, and image reconstruction effects.

The highest-performing MPI tracers have generally been described as monodisperse, single-core magnetite particles with iron core diameters in the range of 20–30 nm [6, 7]. However, a growing number of studies have demonstrated excellent MPI performance from SPIO formulations that would traditionally be considered unconventional or alternative tracers [8–10]. These include large micron-sized iron oxide (MPIO) particles [11–18], multicore particles [6, 19, 20], and chain-like assemblies of magnetic nanoparticles [21–23]. In some cases, these particles exhibit enhanced magnetization dynamics, increased magnetic moments, or altered relaxation properties that improve the MPI signal intensity and/or resolution. Such findings suggest that optimal MPI tracer design may be more complex than initially appreciated and that a broader range of particle architectures may be suitable for certain MPI applications.

For cell tracking applications with MPI, there are additional important considerations in the selection of SPIOs beyond intrinsic magnetic performance alone. First, the SPIO formulation must support efficient and reliable cell labeling. Cell labeling efficiency depends on several factors, including particle size, surface charge, coating chemistry and hydrodynamic diameter [24–28]. Cellular uptake can vary substantially between phagocytic cells such as macrophages, and more difficult to label cells such as lymphocytes. For effective, long-term cell tracking, the goal is for the iron loading per cell to be as high as possible while preserving normal cell function and viability.

Some cell types, and some SPIO formulations, require the use of transfection agents to facilitate or enhance the internalization of particles. Transfection agents promote endocytosis by forming complexes with the particles and altering their interaction with the cell membrane [29]. While this approach improves labeling efficiency, it causes the particles to aggregate which hinders their relaxation resulting in a lower MPI signal [4, 30, 31].

In addition to achieving high cellular iron loading, the intracellular fate of SPIOs is also important for cell tracking with MPI. Following uptake, iron particles are typically compartmentalized within endosomes or lysosomes. This can lead to particle aggregation, immobilization and degradation which can all reduce the MPI signal [30, 32–34]. For example, the peak MPI signal for the highly sensitive SPIO known as Synomag-D is reduced by ∼60% after cell internalization [17]. Therefore, the performance of a SPIO measured in suspension may not accurately predict the MPI performance after cell internalization.

Recent studies suggest that MPIOs may be particularly well suited for robust and quantitative MPI-based cell tracking [11, 12, 14–18]. Importantly, many cell types can be efficiently labeled with MPIOs without the need for transfection agents. In contrast to many conventional SPIO formulations, the MPI signal from MPIO-labeled cells remains comparable to that of MPIOs in suspension, indicating minimal loss of performance following cellular internalization [17]. MPIOs also appear to be relatively resistant to intracellular degradation, which may help preserve MPI signal over time [11]. In addition, labeling with MPIOs typically results in substantially higher intracellular iron loading compared with smaller SPIOs, enabling improved cell detectability [15, 17, 35, 36].

To select the optimal tracers for MPI, researchers are continuing to improve standardized performance metrics. In this study, we characterized and compared the performance of ten commercially available iron particle formulations, spanning a range of magnetic core sizes, hydrodynamic diameters, and surface coatings, using multiple MPR and image-based metrics.

## METHODS

### Superparamagnetic Iron Oxide Particles

Ten commercially available tracers were compared in this study, as outlined in Table 1. VivoTrax (Magnetic Insight Inc.), which is a formulation of ferucarbotran, is a commonly used SPIO for MPI. It consists of aggregated multicore clusters. The core size distribution is bimodal with roughly 70% of small 5 nm cores and 30% of iron cores in the range of 25-30 nm packed together [37]. The entire multicore cluster is enveloped in a carboxydextran coating. VivoTrax+ is fractionated version of VivoTrax which uses magnetic sorting to select for the larger core clusters, resulting in a bimodal distribution with 70% of the particles being 25-30 nm clusters and 30% being 5 nm. The reported hydrodynamic size is 62 nm for both particles; however, independent dynamic light scattering (DLS) data has shown that VivoTrax+ has a significantly larger hydrodynamic diameter (∼90 nm) [38]. Perimag (Micromod GmbH) particles are specifically engineered as a cluster type, or multicore, assemblies with a mean core diameter of 19 nm, a dextran coating and a hydrodynamic size of 130 nm [6]. Synomag-D tracers (Micromod GmbH) are multicore particles with a specialized “nanoflower” core substructure in a dextran matrix that resembles a flower rather than a tightly packed sphere [20]. The mean clustered core size is 30 nm. The nanoflower has a dextran coating. In this study, we compared four Synomag-D particles: two ‘plain’ versions, with hydrodynamic diameters of 50 nm and 70 nm (Synomag-D 50 and Synomag-D 70) and two with a polyethylene glycol (PEG) 25.000-OMe surface modification, with hydrodynamic diameters of 50 nm and 70 nm (Synomag-D PEG 50 and Synomag-D PEG 70). Two MPIOs from Bangs Laboratories were compared, ProMag and Fluorescent Classical Magnetic COOH microsphere (abbreviated to FCM-COOH for this study). Both are microspheres synthesized with a distribution of thousands of iron crystals throughout a polymer matrix with a mean hydrodynamic diameter of 0.9 µm. ProMag is an encapsulated microsphere engineered with an iron-free polymer surface. FCM-COOH is a fluorescent (Flash Red) classical nonencapsulated microsphere, with some iron oxide left exposed on the outer surface. Both versions were functionalized with carboxyl surface chemistry. Feraheme (AMAG Pharmaceuticals Inc.), a ferumoxytol, is an ultra-small SPIO that consists of a single iron core with a mean diameter of ∼7 nm with a specialized semi-synthetic carboxymethyl dextran coating [39, 40]. The mean hydrodynamic diameter is reported to be ∼25 nm.

**Table 1.**
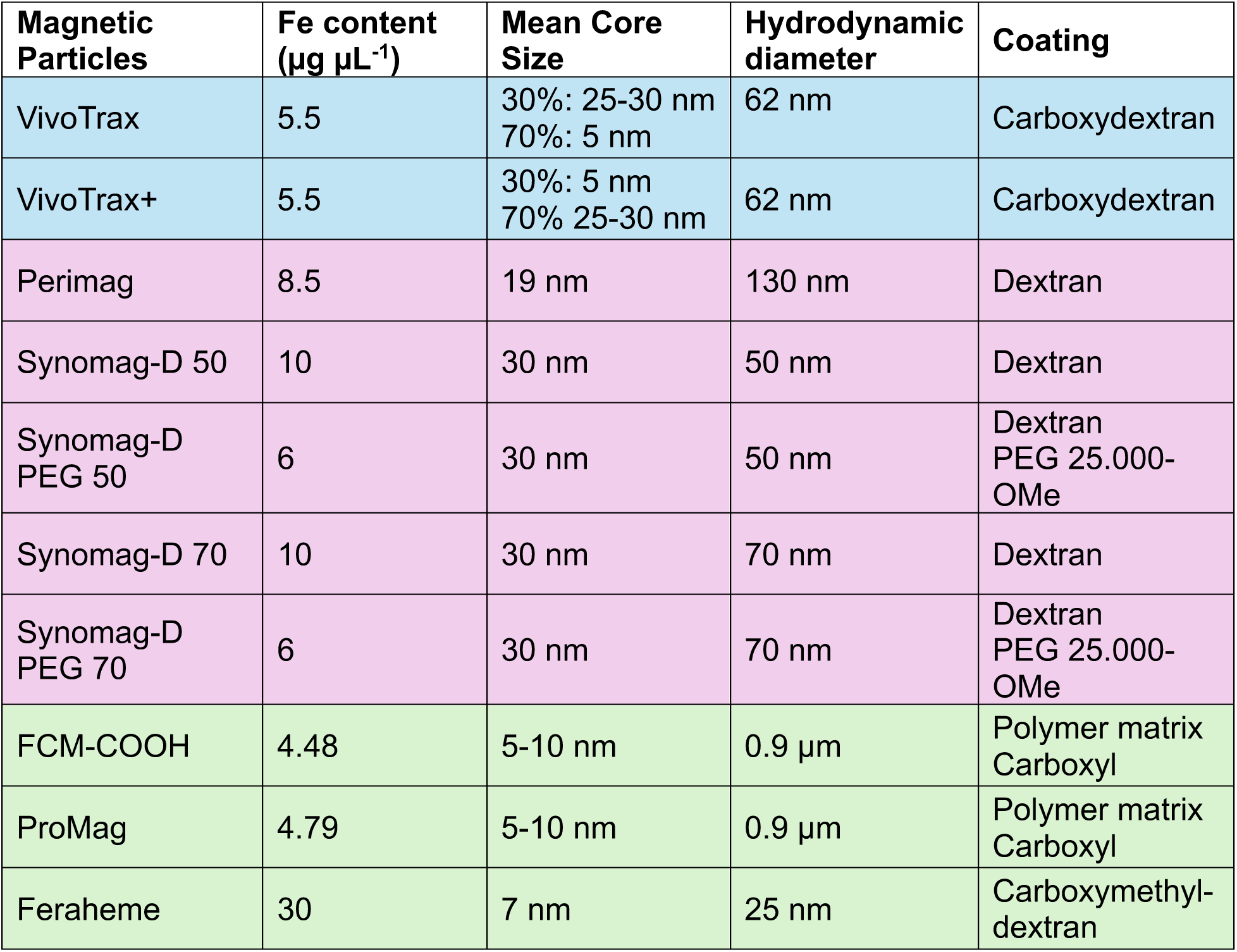
Reported properties of commercially available tracers used in this study.

### Magnetic Particle Relaxometry (MPR)

All tracers were first evaluated using the RELAX module on the MOMENTUM scanner (Magnetic Insight Inc.). Three different sample preparation methods were used to acquire MPR data. In Method 1, a single sample of each tracer was prepared with a volume that resulted in a peak signal intensity of ∼2 A.U. to avoid saturating the receive coil; 2 µL of each tracer was scanned initially, the peak signal was recorded (y), and the volume (x) was adjusted accordingly (2 A.U./y A.U. = x/2 µL). This approach was previously examined in a study where different volumes were compared across two institutions [41]. Method 2 used a single 3 µL sample of each tracer. Method 3 used a single 10 µL sample of each tracer. These volumes were selected because they had previously been reported in the literature [42, 43]. Each sample contained undiluted stock solution in a 0.2 mL PCR tube. Samples were placed vertically in 3D printed holders. Each sample was scanned three times.

### Magnetic Particle Imaging (MPI)

For each tracer, a dilution series, consisting of 5 samples, was prepared within iron mass range of approximately 13 to 220 µg (50 samples in total). 2D images were acquired of each sample individually with a 3 T/m gradient (selection) field and excitation field strengths of 20 and 26 mT in the X and Z channels, respectively. These parameters correspond to the ‘high sensitivity’ imaging mode on the scanner. Before each imaging session, the empty bed with the sample holder was scanned using the same parameters to measure the background signal in the absence of an iron sample.

### Data Analysis

MPR data was analyzed using GraphPad Prism Software (Version 11.0.1) and Excel (Version 16.112.2). The raw peak amplitude of the PSFs, volume and iron mass were measured and recorded for all MPR data. The PSFs were then normalized by iron mass to facilitate comparison between tracers. The peak signal intensity (A.U./µg Fe), also a measure of particle sensitivity, was obtained for all MPR data and the area under the PSF curves were calculated only for MPR data obtained using Method 1. To compare the three different sample preparation methods, the sensitivity (A.U./µg Fe) calculated using Methods 2 or 3 was divided by the sensitivity (A.U./µg Fe) calculated using Method 1.

MPI images were analyzed using open-source HOROS image analysis software (Version 4.0.1). The standard deviation (SD) of the background noise was measured across the entire FOV from images of the empty bed and sample holder. A lower threshold of five times the SD, corresponding to the Rose criterion [44, 45], was used to define the region of interest (ROI) for each image [46]. The area, mean signal, and maximum signal were measured for each ROI. Total MPI signal for each sample was calculated by multiplying the mean signal and ROI area. Calibration lines were created for each tracer from the signals measured for the serial dilution samples by plotting the maximum signal/total MPI signal (y-axis) against the known iron mass of the samples (x-axis). The relationship between MPI signal and iron mass was determined with a simple linear regression, the slope of which represents the amount of MPI signal generated per unit iron mass (A.U./µg Fe); the tracer sensitivity. This line was forced through the origin, under the assumption that, the background MPI signal, in the absence of an iron sample, has an average signal of zero.

MPR data acquired using Method 1 was used when comparing performance metrics between MPR and 2D MPI. The peak signal intensity measured using MPR was compared with slopes of calibration lines created using the maximum signal and the total signal measured from MPI images. The area under the PSF was compared with the calibration slopes created using total MPI signal from images. Simple linear regression and Pearson correlation tests were performed to determine the strength of the relationship between the tracer performance metrics acquired from MPR and MPI.

## RESULTS

Tracer sensitivity measured by MPR is presented in Figure 1 which shows the PSFs obtained for each tracer arranged from highest to lowest peak signal intensity (A.U./µg Fe). The peak signal intensities for the Synomag-D tracers were higher than that of VivoTrax. This agrees with previous studies which have compared these tracers [23, 31, 42]. The peak signal intensity for VivoTrax+ was ∼2x higher than VivoTrax. This agrees with a previous study from our lab [37]. The peak signal intensities for the MPIOs were relatively low. The peak signal intensity for the Synomag-D tracers was ∼ 2x higher than ProMag. The peak signal intensity for ProMag was higher than that for VivoTrax. Feraheme had the lowest peak signal intensity.

**Figure 1.**
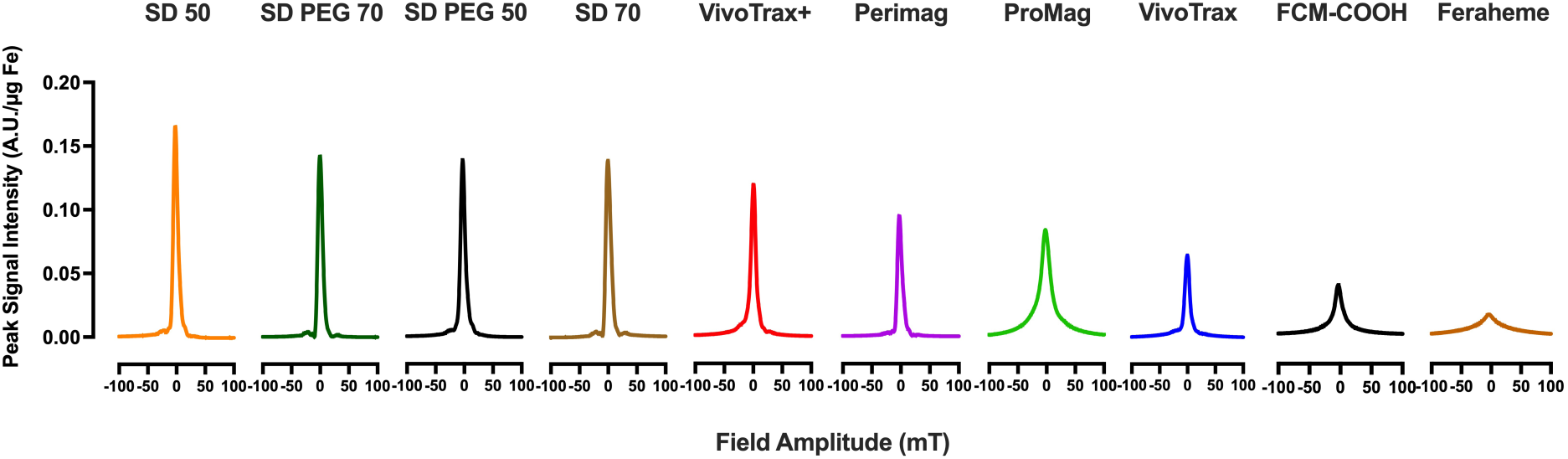
Comparison of tracer sensitivity using MPR. The PSFs obtained for all tracers showing peak signal normalized by iron mass (A.U./µg Fe), arranged from highest to lowest sensitivity, indicated by higher peak signal intensity of the PSF.

Tracer sensitivity evaluated by MPI is shown in Figure 2 which shows the maximum MPI signal per unit iron mass and the calculated slopes of the calibration lines. Note, only three of the samples with the lowest iron masses are included here. This is because when the other two samples with higher iron content were included, the relationship between the maximum signal and iron mass became non-linear due to detector saturation. Tracer sensitivity determined by comparing the maximum signal intensity measured from MPI images was similar to tracer sensitivity reported by MPR. The exception was VivoTrax+ which had a slightly higher slope compared to the Synomag-D PEG tracers. There was a strong correlation between peak MPR signal intensity and maximum signal intensity from imaging (R^2^=0.94; Fig. 3).

**Figure 2.**
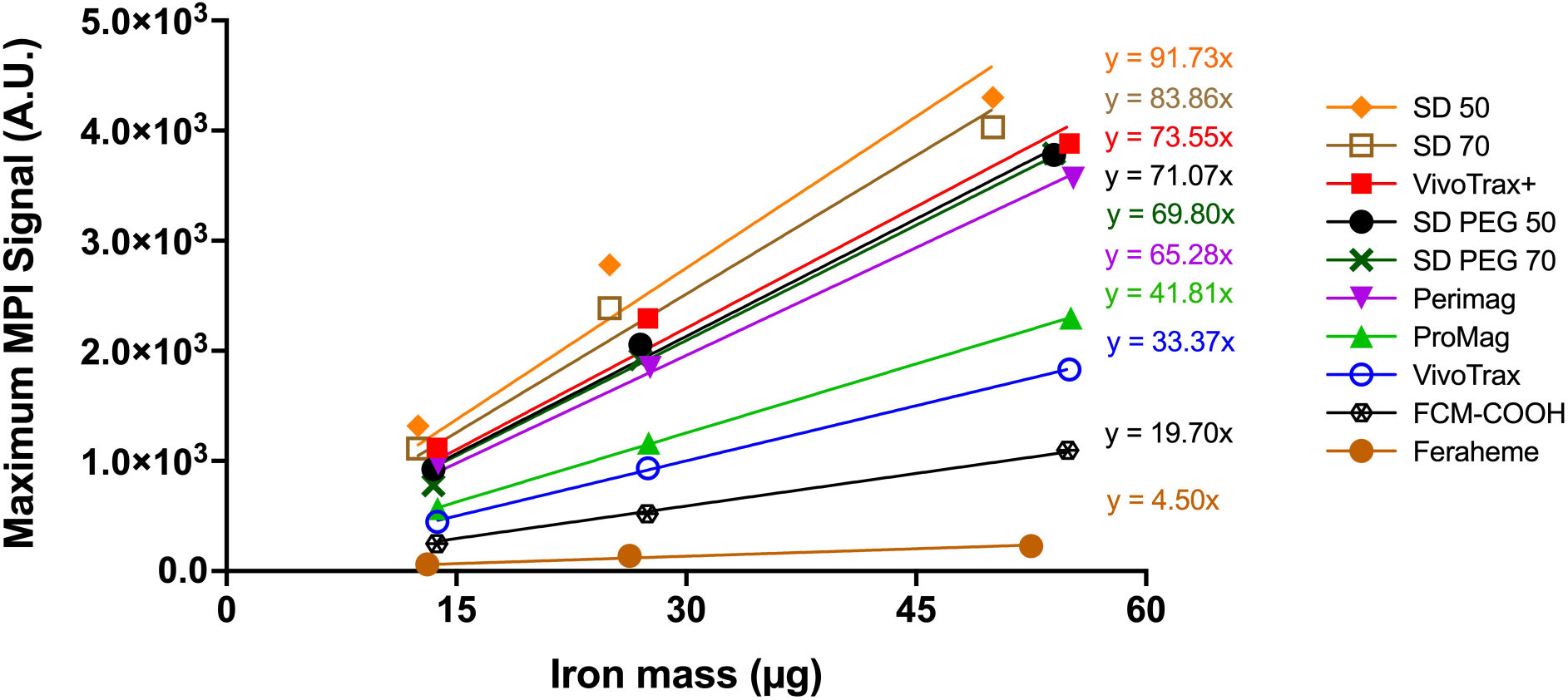
Comparison of tracer performance based on maximum MPI signal measured from 2D MPI images. Maximum MPI signal (A.U.) plotted against known iron mass (µg). The slopes represent the amount of MPI signal generated per µg of iron, referred here as the maximum signal intensity.

**Figure 3.**
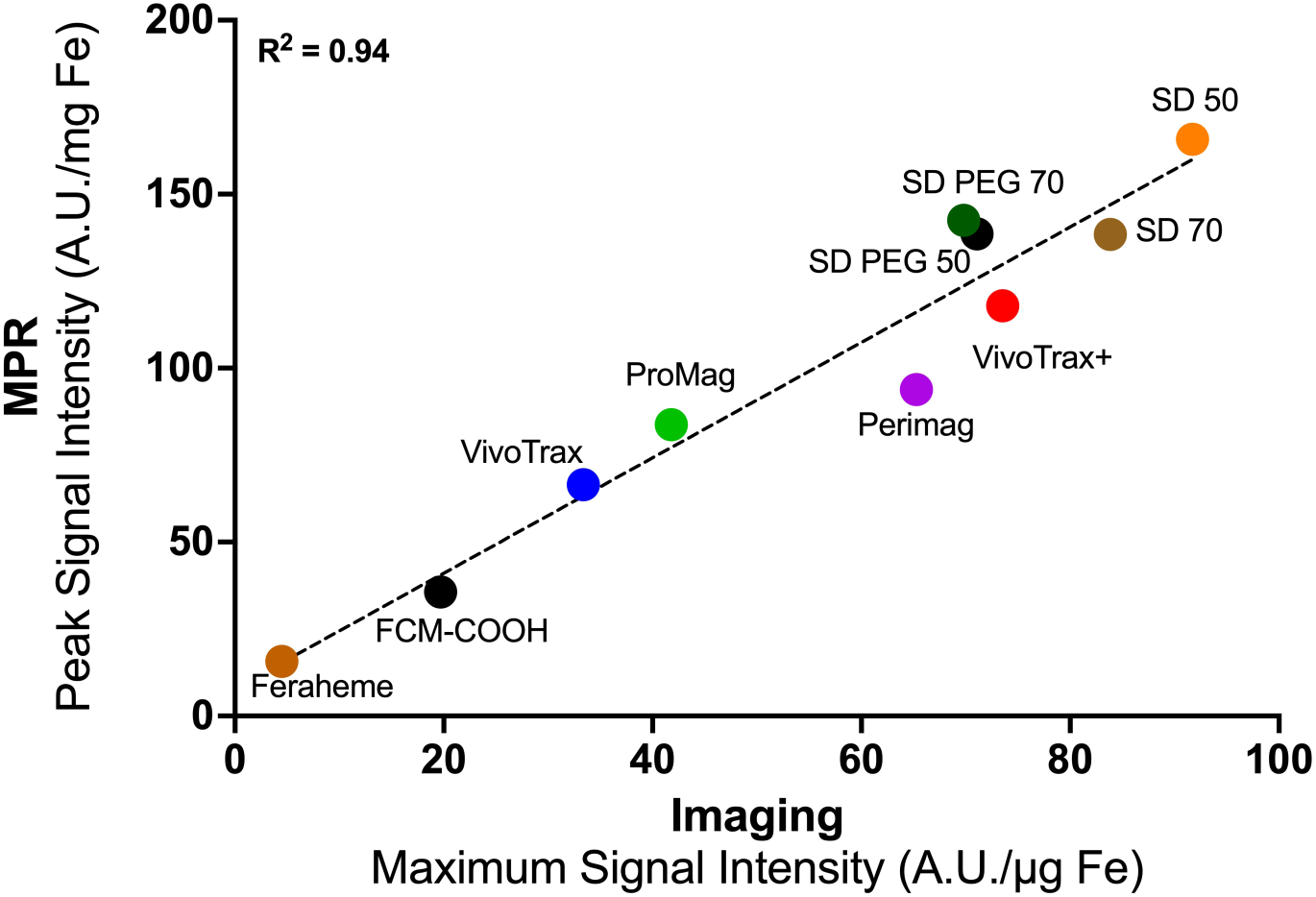
Comparison of peak signal intensity (A.U./µg Fe) from MPR and maximum signal intensity (A.U./µg Fe) measured from 2D MPI images. A strong correlation is observed between the two metrics (R^2^ = 0.94).

Figure 4 shows the total MPI signal per unit iron mass measured from MPI and the calculated slopes of the calibration lines. ProMag had the highest total signal per µg of iron. The slope of ProMag was ∼1.3x higher than Synomag-D 50 and ∼3.2 times higher than VivoTrax. Following ProMag, Synomag-D tracers had the highest total signal per µg of iron with their slopes being ∼1.7-2.5x higher than VivoTrax. The slope for VivoTrax+ was comparable with that for Synomag-D PEG 70. The slope for VivoTrax+ was ∼1.9x higher than VivoTrax. The slope of FCM-COOH was comparable with the slope of Perimag. Both MPIOs, ProMag and FCM-COOH, have higher sensitivity measured by MPI compared to low particle sensitivity measured by MPR (Fig. 5). Interestingly, FCM-COOH, which had one of the lowest peak signal intensities when assessed using MPR, was comparable with Perimag when evaluated using imaging metrics. There was a weak correlation between peak MPR signal intensity and total MPI signal intensity (R^2^=0.38; Fig. 5). There was a strong correlation between the area under the curve for the PSFs and the slopes of calibration lines created using total MPI signal (R^2^=0.89; Fig. 6).

**Figure 4.**
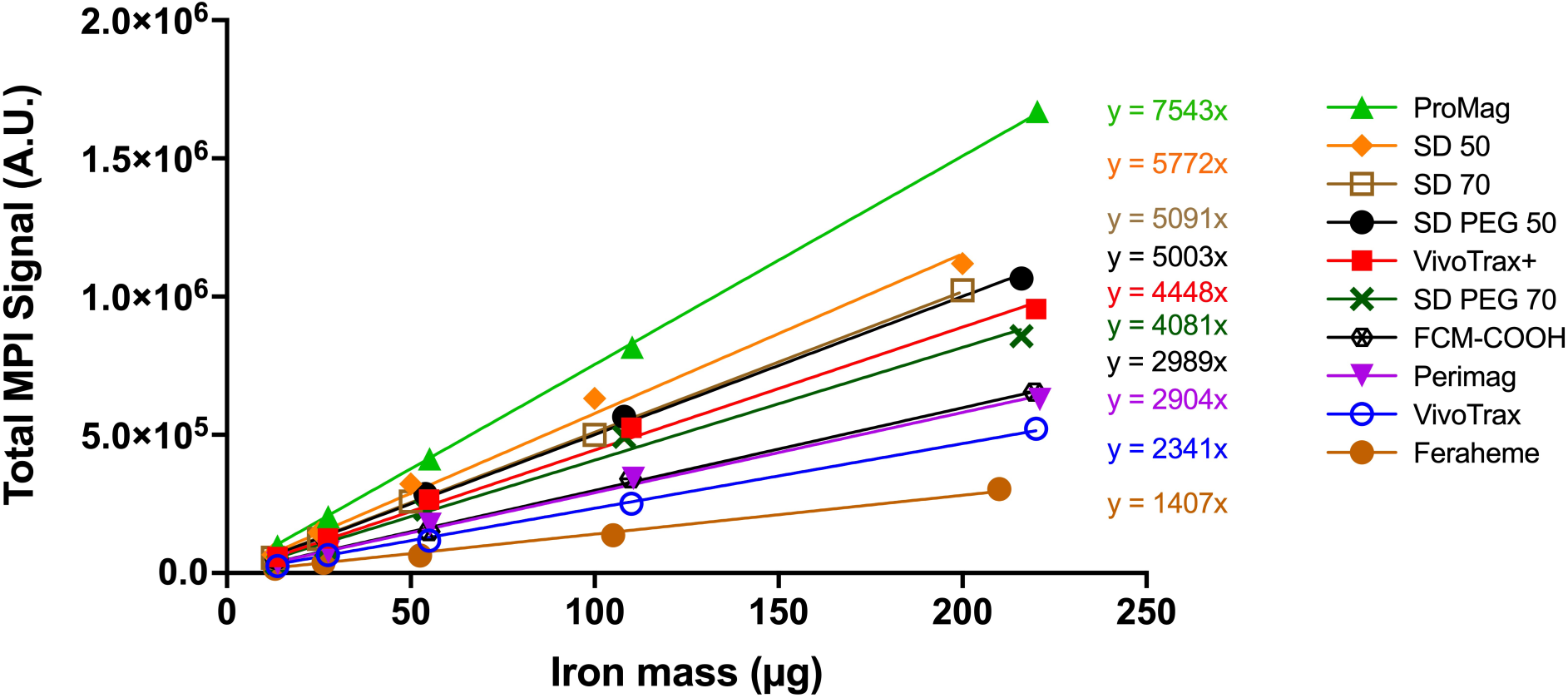
Comparison of tracer performance based on total MPI signal measured using 2D MPI. Total MPI signal (mean signal x ROI area) measured from MPI images plotted against known iron mass. The slopes represent the MPI signal generated per µg of iron, referred here as the total signal intensity.

**Figure 5.**
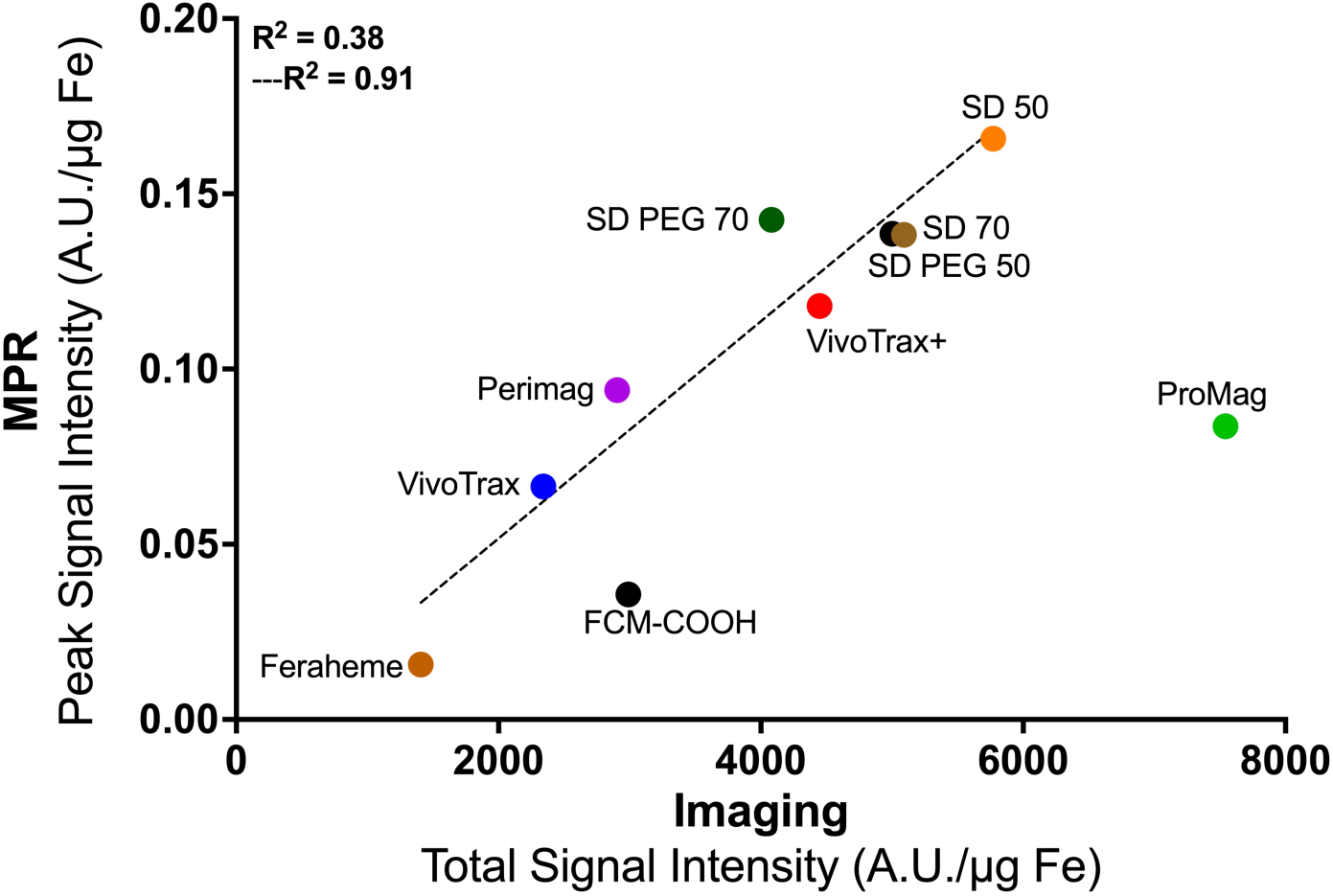
Comparison of MPR peak signal intensity and the slopes of calibration lines created using total MPI signal measured from MPI images. A weak correlation is observed between the two metrics (R^2^ = 0.38). When both MPIOs (ProMag and FCM-COOH) are excluded from analysis, the correlation improves (---R^2^ = 0.91; dashed line).

**Figure 6.**
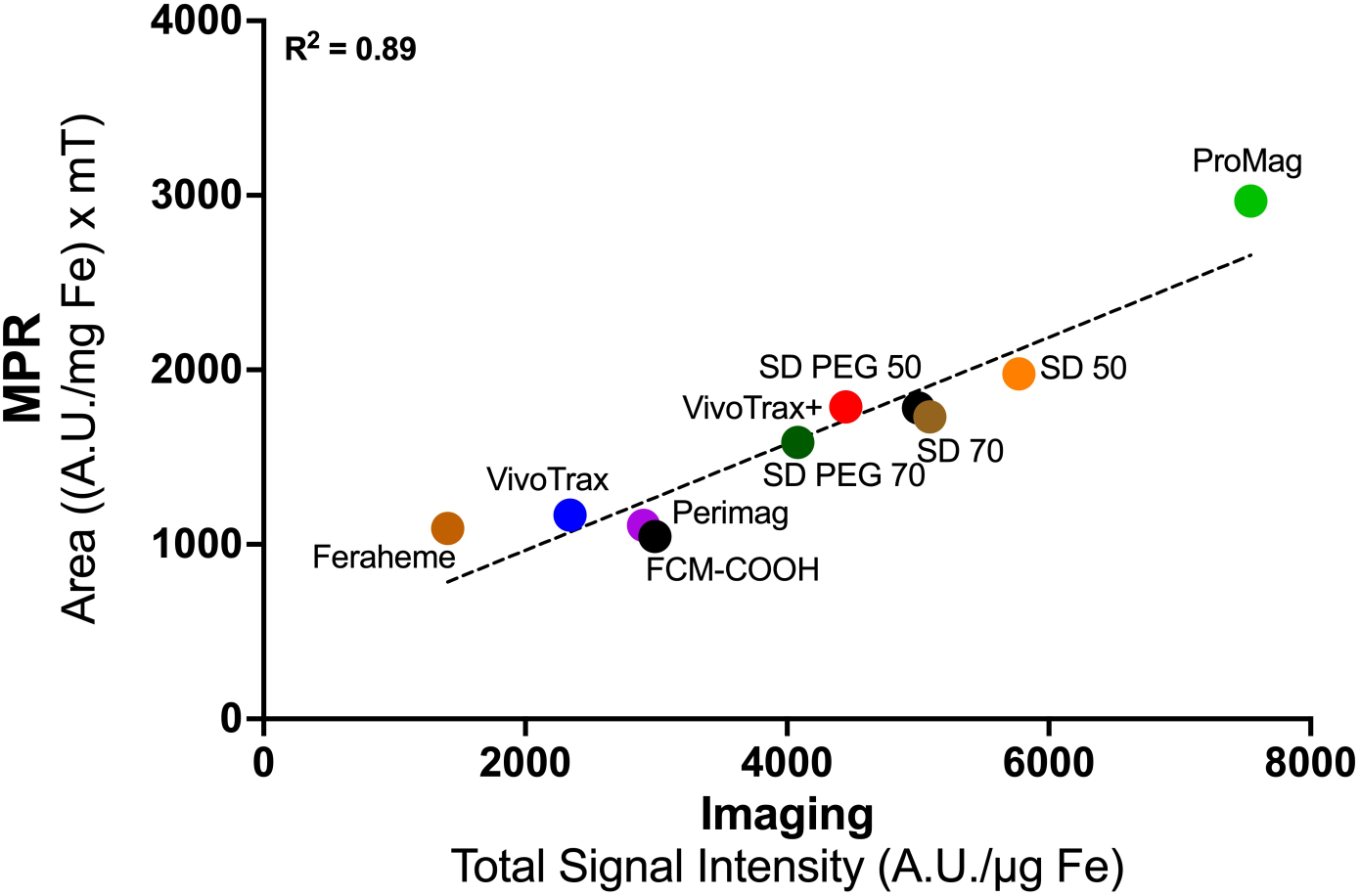
Comparison of area under the PSF measured using MPR and the slopes of calibration lines created using total MPI signal from 2D MPI images. A strong correlation is observed between the two metrics (R^2^ = 0.89).

Figure 7 shows the results of MPR data acquired using variable sample volumes, optimized to produce a target raw peak voltage amplitude of ∼ 2 A.U., and fixed sample volumes of 3 and 10 µL. Table in 7A shows the sample volume and iron mass along with the amplitude (A.U.) and the calculated particle sensitivity (A.U./µg Fe). Figure 7B shows sensitivity measured using Methods 2 and 3 relative to Method 1. Bars under the 100% line indicate reduced sensitivity compared to Method 1. For all tracers, except for VivoTrax, the sensitivity determined using Method 2 was close to that measured for Method 1. In contrast, Method 3 resulted in lower sensitivity for most tracers with Synomag-D and Perimag tracers having the most notable reduction. Figure 7C quantifies the non-linear response by comparing the change in iron mass and amplitude for Methods 2 and 3 relative to Method 1. For Synomag-D tracers and Perimag, the increase in iron mass did not yield a proportional increase in amplitude, with ∼30-58% signal lost compared to Method 1.

**Figure 7.**
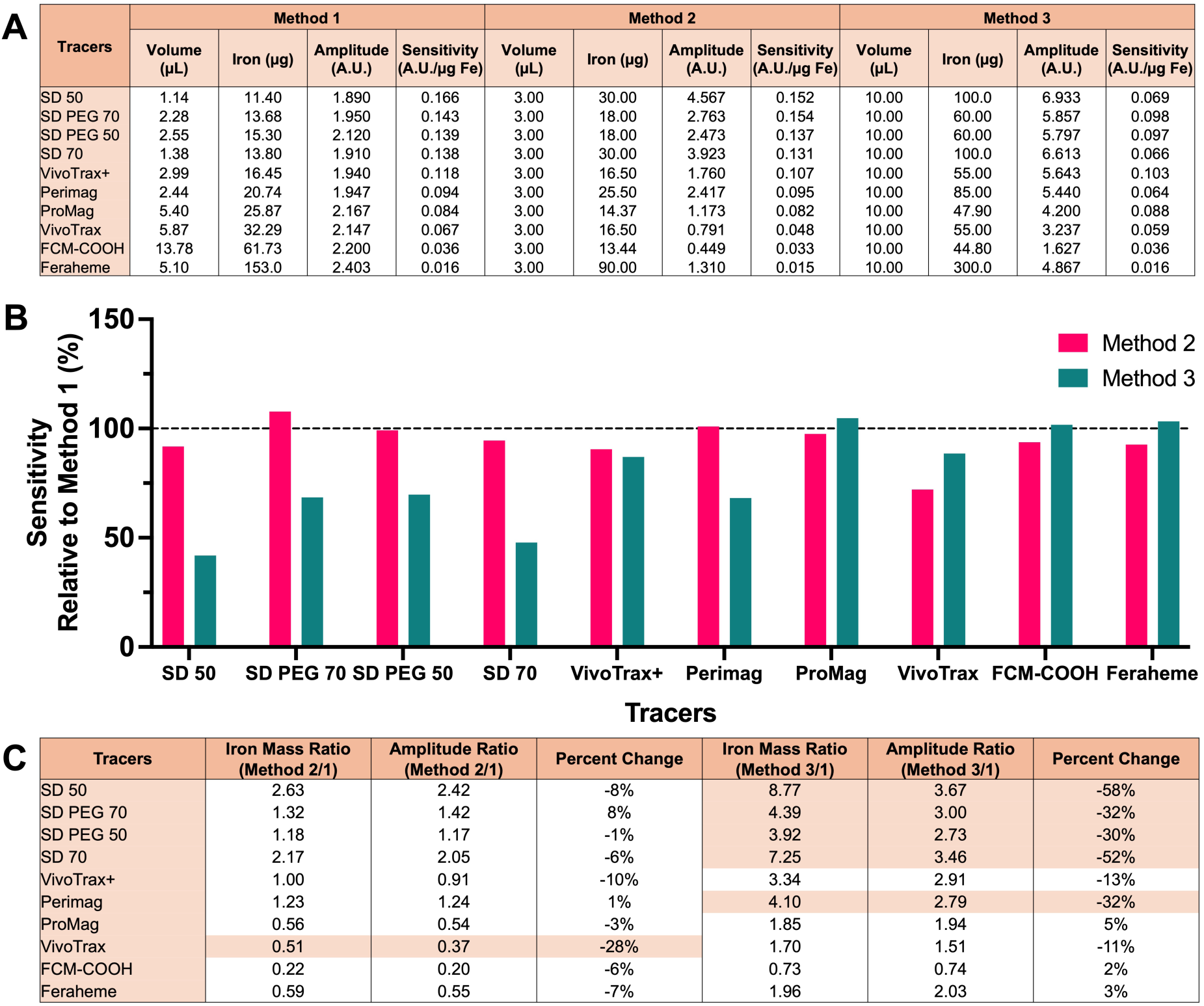
Comparison of tracer sensitivity measured by MPR using three different sample preparation methods. (A) The table shows the sample volume, iron mass, the amplitude and the calculated sensitivity for each SPIO. (B) The tracer sensitivity calculated for Methods 2 and 3 relative to Method 1. (C) The table shows the ratio of iron mass, the ratio of the amplitude and the percent change for samples prepared using Methods 2 and 3 relative to Method 1. The orange shaded cells highlight samples that suffered a signal deficit greater than 25%.

## DISCUSSION

In this study, we characterized and compared ten commercially available tracers using MPR and MPI-based quantitative metrics. Synomag-D tracers showed the highest peak signal intensities when measured using MPR and the highest maximum signal intensity in 2D MPI. Synomag-D tracers were synthesized specifically for MPI as a cohesive 30 nm multicore ‘nanoflower’ cluster. Its physical structure consists of sub-cores, or ‘petals’ which are ∼5 nm. These are fused together and have matching orientations which leads to very strong coupling. Structurally, the entire 30 nm nanoflower behaves like a single crystal and the core size matches the theoretical Langevin maximum magnetic power. While a standard single core 30 nm particle uses Brownian relaxation and encounters phase lag when responding to the rapid magnetic field changes in MPI, for Synomag-D, the magnetic moments flip mainly via Neel relaxation leading to a higher voltage spike. For standard particles, the speed and power cannot be changed independently, they are locked together by the particles size. Synomag-D particles defy this rule; by fusing 5 nm petals into a 30 nm flower, it creates a structural layout which produces a signal stronger than small Neel particles and large Brownian particles. Our results align with other studies which have compared Synomag-D to commercial or custom engineered MPI tracers [23, 42].

The 50 nm hydrodynamic size variant of Synomag-D produced a higher peak MPR signal and higher maximum and total MPI signal compared to the 70 nm variant. This agrees with Kumar et al. who compared both to nanochains developed for MPI [23]. Even though both particles have the same 30 nm clustered core size, the 50 nm particle has a higher ratio of magnetic core to polymer shell allowing for tighter magnetic coupling and boosting the collective magnetic moment. In addition, although Synomag-D is Neel dominant, a minor fraction of its magnetic behaviour relies on Brownian relaxation which scales cubically with the hydrodynamic radius. Increasing the hydrodynamic diameter from 50 to 70 nm almost triples the hydrodynamic volume and this increases the frictional drag reducing peak signal intensity for the larger particle.

The functionalized Synomag-D tracers exhibited a lower maximum and total signal intensity in MPI compared to plain Synomag-D. This is known to occur because surface modification alters the magnetic core packing and shields the dipole-dipole interactions. This was also the case when comparing the peak signal intensity for Synomag-D 50 and Synomag-D PEG 50 measured by MPR. Although, no difference in the peak signal intensity was observed by MPR for the 70 nm variants.

We evaluated two versions of the multicore tracer VivoTrax. VivoTrax is considered a polydisperse, bimodal aggregate. The physical structure of VivoTrax also consists of small iron cores which are ∼5 nm. However, unlike Synomag-D tracers, clusters are formed randomly by natural or spontaneous aggregation of the small cores into clumps.

The result is a wide mix of cluster sizes and geometries. Approximately 70% of the iron mass consists of isolated 5 nm cores or tiny, weak aggregates lacking the magnetic volume to contribute to the MPI signal. As expected, VivoTrax+ had higher signal compared to VivoTrax for all metrics. This is because it undergoes a magnetic fractionization process which optimizes the core size distribution. We reported this in a previous study along with demonstrating that VivoTrax+ shows enhanced cellular internalization [37]. Even though VivoTrax+ is enriched with more of the larger 25-30 nm cores, the MPI signal is lower compared to Synomag-D. This is likely because the larger clusters are loosely bound, chaotic aggregates of 5 nm sub-cores with imperfect coupling, rather than an optimized nanoflower structure. In addition, VivoTrax+ still retains about 30% of the very small 5 nm cores, which act as non-contributing ‘dead weight’.

The MPI signal intensity for Perimag was lower than the Synomag-D tracers and VivoTrax+, but higher than VivoTrax, for all metrics, which agrees with the findings by Imhoff et al [42]. Like VivoTrax, Perimag is categorized as a multicore particle. However, Perimag maximizes the fraction of iron that falls within the optimal range for MPI by packing very small (3-8 nm) crystals tightly into dense clusters which are ∼19 nm [6]. This is the main reason why Perimag signal is higher than VivoTrax. The hydrodynamic size of Perimag is much larger than Synomag-D and VivoTrax+ (130 nm versus 60-70 nm). The larger size creates more fluid friction and physical drag compared to Synomag-D and VivoTrax+ that decreases the MPI signal intensity. Feraheme produced the lowest MPI signal measured by all metrics. This is because it is an ultra-small superparamagnetic iron oxide with a very small iron core cluster diameter (∼7 nm) which translates to a small magnetic moment and a weak MPI signal.

We evaluated two types of MPIOs from Bangs Laboratories, ProMag and FCM-COOH. Both had a relatively low peak signal intensity when measured using MPR and a low maximum signal intensity measured from 2D MPI. Both MPIOs have a massive hydrodynamic diameter which experiences immense viscous friction, such that the physical rotation (Brownian relaxation) is effectively zero. Any magnetic response from the MPIOs is driven by internal Neel relaxation. The low peak signal intensity is influenced primarily by the tight spacing of thousands of small cores into the rigid polymer matrix which causes very strong magnetic dipole-dipole coupling. This substantially delays the internal Neel relaxation dragging out the magnetic response and flattening the voltage peak. The peak signal intensity for ProMag was approximately double that for the FCM-COOH. This is likely because FCM-COOH has a higher iron concentration (∼60%) with the iron crystals crowded more closely together. ProMag has a lower iron concentration (∼20-25%) and the cores are more evenly distributed enabling them to flip more freely. In addition, ProMag are described as geometrically perfect, monodisperse spheres while FCM-COOH has irregular shapes with a much wider size distribution profile (0.2 to 1.5 µm). Because of the polydispersity of FCM-COOH, the magnetic response is asynchronous with different sized particles responding differently to the drive field leading to a weaker signal compared to ProMag particles for which the magnetic flipping is more synchronous.

When comparing MPR and MPI metrics for evaluating tracer sensitivity, there was a strong correlation between the peak signal intensity measured by MPR and the maximum signal intensity measured by MPI (R^2^ = 0.94). There was also a strong correlation between the area under the curve measured by MPR and the total signal intensity measured by MPI (R^2^ = 0.89). Overall, MPR measurements can be used to predict MPI performance for most tracers. This has been confirmed in several previous studies [23, 42].

While MPR is an essential tool to determine the intrinsic sensitivity of tracers, it does not completely replicate the imaging conditions. The gradient and drive fields used for MPI and the physical environment of the tracer can alter how signals are generated, and thresholding and reconstruction methods can impact the amount of signal measured from images. Our results showed that ProMag had a relatively low peak MPI signal intensity measured by MPR, but the highest total MPI signal per µg of iron measured from images. The total MPI signal measured from images reflects the integrated signal across a reconstructed ROI and is calculated by multiplying the mean signal by the area of the ROI. When comparing the metrics from images of ProMag and Synomag-D, we observed that the area of the ROI was similar, but the mean signal intensity was higher for ProMag resulting in a higher total MPI signal (Fig. S1). This can be understood by considering the PSF shape, how the signal is distributed across pixels, and the 5 x SD threshold. Synomag-D creates a sharp, tall, narrow PSF. Only a small cluster of pixels at the center of the ROI capture the very high maximum signal. The surrounding pixels immediately drop off into an area of very low-amplitude phase-lag tails. With a 5 x SD threshold, the mask expands to capture those low signals in the outer tails. Averaging the smaller number of very high center voxels with the much larger number of low tail voxels pulls down the calculated mean for Synomag-D. Although the PSF for ProMag exhibits a lower maximum signal intensity, its mean signal value within the 5 x SD mask surpasses that of Synomag-D because the thresholded ROI encompasses the low-amplitude peripheral signals.

Measuring the total MPI signal from images is relevant because the total integrated signal is linear with the amount of iron tracer present. Importantly, the iron mass can only be calculated using the total MPI signal and a calibration line (Iron mass (y) = mx; where m is the slope of the calibration line and x is the total MPI signal from the image ROI). A higher total MPI signal translates to superior sensitivity for cell tracking and quantification. In general, one should consider that the PSF measured by MPR evaluates a tracer at its theoretical best, whereas an MPI image captures the tracer’s actual performance of a tracer under imaging conditions.

In addition to generating a higher total signal, MPIOs offer distinct advantages over SPIOs for *in vivo* cell tracking with MPI. Small nanoparticles like VivoTrax or Synomag-D often suffer from a reduction in MPI signal once internalized by cells due to aggregation and constrained rotation inside endosomes/lysosomes [30, 47]. Larger polymer-encapsulated particles like ProMag maintain a stable MPI signal that does not significantly differ between free and intracellular states [17]. Achieving a high mass of iron per cell with SPIOs can require the use of transfection agents, which can alter cell viability and reduce the MPI signal due to aggregation [30, 37]. MPIOs contain much more iron per particle compared to SPIOs and cells can be easily labeled without transfection resulting in higher iron loading of cells. SPIOs are quickly broken down by intracellular lysosomal enzymes, causing a rapid loss of the tracer signal over time [47]. The polymer shell surrounding the iron cores in MPIOs protects them from rapid enzymatic degradation, facilitating long-term longitudinal tracking of cells [35]. There are some practical limitations for using MPIOs for MPI cell tracking. MPI resolution of MPIOs is much lower (∼6-7 mm) compared to SPIOs (∼2-3 mm; Table S1). Additionally, because they are not degraded, MPIOs persist *in vivo* restricting their use to preclinical research. While only minor changes in cell phenotype or function are reported for cells labeled with MPIOs, studies have shown that MPIOs can impair stem cell differentiation, and immune cell maturation and migration [48, 49]. The trade-offs for cell tracking with SPIOs and MPIOs is summarized in Table S2.

To prepare samples for MPR, we use variable volumes optimized to produce a target peak voltage amplitude of ∼ 2 A.U. (Method 1), which represents a compromise between maximizing the system signal to noise ratio and maintaining a safety margin on detector saturation. At approximately 5 A.U., the system is at maximum voltage capacity. One cause of detector saturation is the mass of iron, which is dictated by the sample volume. For some tracers, a very high mass of iron can cause a voltage spike that overloads the detector. This results in distortion or clipping of the peak of the raw signal amplitude which subsequently decreases the peak signal intensity of the processed PSF. In a non-saturated system, a 2x increase in iron mass should yield a 2x increase in peak signal. Our data showed that for the 10 µL samples (Method 3) of the Synomag-D tracers and Perimag, the change in peak signal did not keep up with the change in iron mass indicating some detector saturation under these conditions. For example, for Synomag-D 50, the iron mass increased by 8.8x between Methods 1 and 3 but the signal only increased by 3.6x; indicating that the detector only recorded ∼ 40% of the true magnetic response from the 10 µL sample. The peak voltage amplitudes for these samples were between 5.4 and 6.9 A.U. As a result, the tracer sensitivity (A.U./µg Fe) calculated for these 10 µL samples is inaccurate and artificially low.

The magnetic properties (core size, saturation magnetization and relaxation dynamics) of the tracers also play a large role in detector saturation, acting as a multiplier on the iron mass. A highly sensitive tracer like Synomag-D effectively amplifies the voltage response per unit mass, lowering the threshold for detector saturation. Our data show that the high-performance tracers, like Synomag-D, require a lower iron mass to hit the 2 A.U. target. For magnetically inefficient tracers, even very high iron masses do not lead to detector saturation. While a 10 µL sample of Synomag-D 50 (100 µg Fe) generates a sharp voltage spike that saturates the detector at 6.9 A.U., a 10 µL sample of Feraheme with 3x the iron mass (300 µg Fe) peaks safely at 4.9 A.U. without saturating the electronics. This is because Feraheme has a very broad PSF that scales smoothly without producing the sharp voltage spikes that overwhelm the receiver electronics.

There are some experimental details which could be considered limitations of this study. When we compared the MPR data acquired using different sample preparation methods, we only used one sample per tracer so we could not report statistics on differences between tracers. We, and others, have previously found no significant difference in the peak MPI signal (A.U./µg Fe) measured by MPR across three independent samples [42]. Therefore, a single sample was prepared using the variable volume method and 3 repeated measurements were acquired per sample. For the imaging data we only used one acquisition mode (called 2D high sensitivity mode on the Momentum scanner). The use of different gradient field and drive field strengths will affect the MPI signal of the tracers.

## CONCLUSION

In this study, we characterized and compared ten commercially available tracers with various magnetic properties using MPR and image-based quantitative metrics. We found that Synomag-D tracers have the highest peak signal intensity using MPR and maximum signal using imaging, while ProMag, a MPIO, had the highest total signal intensity when evaluated using imaging. ProMag has the potential for *in vivo* preclinical MPI cell tracking applications due to its size which can facilitate high iron loading and high total signal intensity which is used for robust cell quantification. We also show that while MPR can estimate the theoretical MPI performance of tracers, a comprehensive evaluation using imaging is needed to fully assess the potential of MPI tracers. This work offers a thorough comparison for careful tracer selection including considerations for future preclinical MPI cell tracking studies.

---

**Figure S1.**
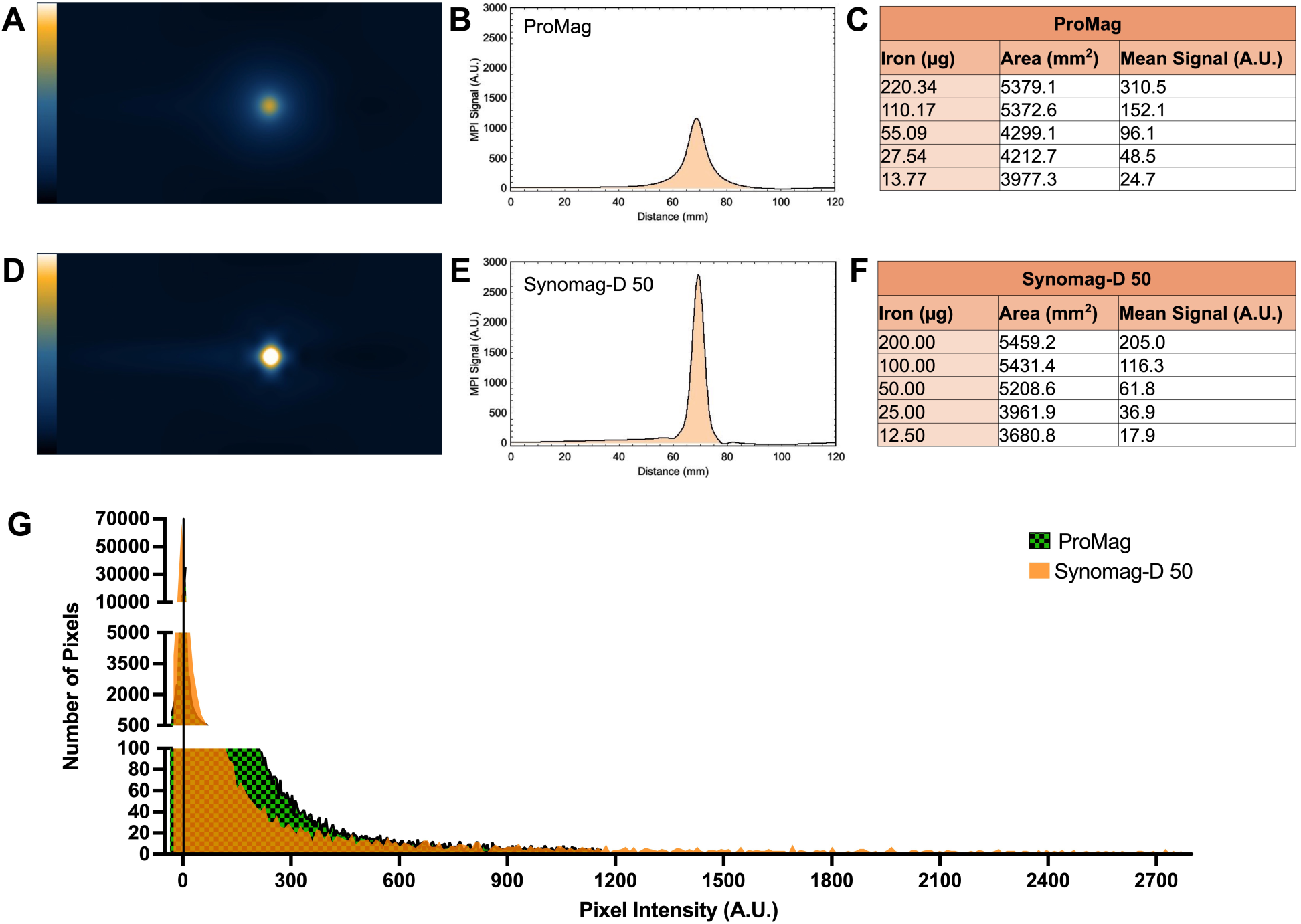
Comparison of metrics measured from MPI images for ProMag and Synomag-D 50. Representative MPI images, adjusted to the same colour scale (-25 to 3000 A.U.) for visual comparison for (A) ProMag and (D) Synomag-D 50. Line profiles were created for (B) ProMag and (E) Synomag-D 50 across the z-direction of the MPI image. The area of the ROI and the mean signal measured from images of the dilution series with 5 samples, using 5 x SD threshold method for ROI segmentation, for (C) ProMag and (F) Synomag-D 50. (G) Pixel intensity distribution histogram for a representative ProMag and Synomag-D 50, illustrating the relative proportion of mid-range signal intensity pixels with ProMag compared to Synomag-D 50. The solid black line at x = 1.8 represents the lower threshold for ROI segmentation when using the 5 x SD analysis method. Line profiles were created using Fiji, an image analysis software that is a “batteries-included” distribution of ImageJ (https://imagej.net/software/imagej).

**Table S1.** Full-width half maximum (FWHM; mT) and resolution (mm) measured using MPR. System reported FWHM values were divided by gradient strength of 3 T/m to acquire resolution in millimetres. Tracers are ordered from highest to lowest resolution.

| Particles | FWHM (mT) | Resolution (mm) |
| --- | --- | --- |
| SD PEG 70 | 8.28 | 2.76 |
| SD 50 | 8.38 | 2.79 |
| Perimag | 8.61 | 2.87 |
| SD PEG 50 | 8.92 | 2.97 |
| SD 70 | 8.96 | 2.99 |
| VivoTrax+ | 9.62 | 3.21 |
| VivoTrax | 9.79 | 3.26 |
| FCM-COOH | 19.26 | 6.42 |
| ProMag | 23.33 | 7.78 |
| Feraheme | 56.72 | 18.91 |

**Table S2.** Summary of key characteristics of SPIOs and MPIOs to consider for cell tracking applications with MPI.

| Characteristics | SPIOs | MPIOs |
| --- | --- | --- |
| Typical physical structure | Single core or multi-core structure with aggregates of iron cores | Large polymer matrix containing thousands of iron cores embedded within |
| Typical hydrodynamic diameter | in nanometers range (~20-150 nm) | in micrometers range (~0.93-1.63 $\mu\text{m}$) |
| Iron content per particle | low | high (~1 pg Fe per particle) [18, 50] |
| Magnetic saturation behaviour | Steeper M-H curves | Broad M-H curves |
| Resolution | High resolution indicated by narrow PSF (~3 mm) | Low resulting in slower relaxation and broadening of PSF (~7-8 mm) |
| Cell labeling efficiency | Requires transfection agents to achieve sufficient iron loading [15, 31, 51] | High cellular iron loading with no transfection agents required [15, 35] |
| Intracellular signal stability | Decrease in MPI signal following cellular internalization [11, 30, 47] | Similar MPI signal before and after cellular internalization [17] |
| Common examples of commercially available tracers | VivoTrax, Synomag-D | ProMag, FCM-COOH |

## Notes

### Competing Interest Statement

The authors have declared no competing interest.

### Funding

- Canadian Institutes of Health Research
- Natural Sciences and Engineering Research Council

